# Genome-wide identification of the oat *MLO* family and identification of a candidate *AsMLO* associated with powdery mildew susceptibility

**DOI:** 10.1101/2021.03.22.435145

**Authors:** Aisling Reilly, Hesham A.Y. Gibriel, Sujit Jung Karki, Anthony Twamley, John Finnan, Steven Kildea, Angela Feechan

## Abstract

*Blumeria graminis* f. spp. *avenae* is the causal agent of powdery mildew disease in oats (*Avena sativa*). It is the most significant limiting factor to oat production, with yield losses ranging from 5%-40%, during high disease pressure conditions. Certain members of the *Mildew Locus O* (*MLO*) gene family have been shown to act as powdery mildew susceptibility factors in many different plant species. A loss-of-function mutation of specific *MLO* genes confers broad-spectrum resistance against powdery mildew pathogens. Potential *MLO* candidates have not yet been identified in oats. In this study, we identified oat MLOs by querying 341 known MLO protein sequences against the publicly available oat genome. 11 MLO-like sequences were identified in oats. Phylogenetic analysis grouped these candidates into four different clades, one of which, AsMLO1 was grouped together with other cereal MLOs functionally known to contribute to powdery mildew susceptibility. AsMLO1 showed the highest similarity to the known powdery mildew-associated MLO proteins from wheat and barley. Gene expression analysis revealed *AsMLO1* expression is up regulated at 12 hours post-infection with *Bga* and was inferred to be a candidate gene associated with powdery mildew susceptibility in oats. These results are an important step towards more durable strategies to control powdery mildew incidence and severity in oats.

## Introduction

The *Mildew Locus O* (*MLO*) gene family are a plant-specific family of genes that have been identified in many agronomically important crop species (Iovieno *et al.*, 2015). While the biological function of these proteins remains unknown (Kusch, *et al.*, 2016), some *MLO* gene members have been shown to be negative regulators of plant defence responses against powdery mildew pathogens (Cui *et al.*, 2018), while other *MLO*s have been associated with developmental process, such as pollen tube elongation and root development (Chen *et al.*, 2020).

The powdery mildew-associated *MLO* was first identified when a natural loss–of-function mutation occurred in barley, resulting in broad-spectrum resistance against *B. graminis hordei* (*Bgh*) (Kusch and Panstruga, 2017; Jørgensen, 1992). Since its initial discovery in the 1930s (Jørgensen, 1992), *mlo*-mediated resistance has been widely used in spring barley cultivars across Europe and has remained effective since its deployment (Pavan *et al.*, 2010; McGrann *et al.*, 2014). Subsequently, many orthologues of the powdery mildew-responsive *HvMLO* have been identified in other agronomically important crop species, such as wheat (Acevedo-Garcia *et al.*, 2017), rice (Nguyen *et al.*, 2016) and tomato (Appiano *et al.*, 2015), to name but a few.

During the early stages of powdery mildew infection, *MLO* becomes localised at the site of fungal penetration and negatively affects the components of two penetration (PEN) resistance pathways, PEN1/ROR2 and PEN2/PEN3 defence pathways (Underwood and Somerville, 2008). The mechanisms by which powdery mildew pathogens manipulate *MLO* expression to promote fungal infection and negatively regulate plant defences remains largely unknown, although some studies have suggested that pathogen effectors may play a role (Niks, *et al.*, 2015).

*MLO* genes encode proteins that are characterized by the presence of seven transmembrane (TM) domains, a calmodulin binding domain (CaMBD) at the C-terminus and an extracellular N-terminus (Kusch and Panstruga, 2017). These features result in the formation of three extracellular and three cytoplasmic topological loops (Chen *et al.*, 2020) and are highly conserved in MLOs from different plant species (Kusch, *et al.*, 2016). Extensive phylogenetic analyses shows that MLO proteins are grouped into subfamilies of up to eight well-defined clades (Rispail and Rubiales, 2016) (I to VIII) and are typically grouped depending on individual MLO function (Chen *et al.*, 2020). For example, disease-associated MLOs from monocot and dicot plant species are found within Clade IV and Clade V, respectively (Elliott *et al.*, 2002; Kusch, *et al.*, 2016). The highly conserved features of the MLO family suggests that tools such as phylogenetic analyses, multiple alignments and gene expression analyses can be used to identify new MLO homologs associated with powdery mildew susceptibility in many plant species (Iovieno *et al.*, 2015).

Oats (*Avena sativa*) is one of the major cereal crops grown worldwide and is ranked as the sixth most important cereal crop globally (Isidro-Sánchez *et al.*, 2020). The significant limiting factor to oat production is the incidence and severity of fungal pathogens, such as powdery mildew (Clifford, 1995) caused by the biotrophic fungal pathogen *Blumeria graminis* f. spp. *avenae* (*Bga*) (Sánchez-Martín *et al.*, 2016). Due to its infectious life cycle, *Bga* can be difficult to control and eradicate in a crop once established (Roderick, *et al.*, 2000). Currently, the most sustainable method of controlling *Bga* in oats is the use of resistant cultivars (Okoń, 2015). However, only 11 resistance genes have been catalogued in oats (Ociepa *et al.*, 2020), and the effectiveness of these *R-*genes may break down over time with the emergence of new virulent isolates of the pathogen (Várallyay, *et al.*, 2012). As a result, there is a need to identify more durable forms of resistance. Due to the longevity already demonstrated by *mlo*-mediated resistance in barley, it offers a unique opportunity to provide such durable resistance in oats.

Using the recently available oat genome sequence (Blake, *et al.*, 2019), this study identified the *MLO* gene family and determined a potential candidate acting as a powdery mildew susceptibility factor in oats.

## Materials and Methods

### Plant Materials and Powdery Mildew Inoculations

The *Bga*-susceptible oat cultivar Barra was used in this study. Seeds were stratified for three days at 4°C and then incubated in darkness at room temperature for germination. Germinated seeds were then placed in pots containing John Innes No. 2 compost. Plants were grown in chambers with 16 hours light at 22°C and 8 hours darkness at 16°C until three weeks old.

A single spore isolate of *Bga* was previously collected and cultured from a field isolate found at UCD Lyons Research farm, Co. Kildare. This single spore isolate was maintained on Barra plants in a containment unit under natural light and temperature. For inoculations, oat seedling plants were placed at the bottom of a settling tower. Up to three *Bga-*infected oat plants were used as a source of inoculum by gently shaking them over the settling tower, as outlined in Twamley, *et al.*, (2019). Plants in the tower were left for 10 minutes to allow the spores to settle.

### Microscopy

Detached leaves were subsequently stained by heating the leaves in Trypan blue solution for 20 minutes (Koch and Slusarenko, 1990). Leaves were then destained in chloral hydrate solution (2.5g/ml H20) overnight. Samples were mounted onto microscope slides with 80% glycerol. To establish a time-course of *Bga* infection on oats, leaves infected with *Bga* were collected at 0, 8, 12, 24, 48 and 72 hours post-inoculation and stained using trypan blue method (Feechan *et al*., 2009). Slides were examined under bright field light using a Leica DM5500B microscope and images were taken with a Leica DFC310FX digital camera.

### Identification of *AsMLO1*

Prior to the publication of the *Avena sativa* genome, an *Avena barbata* wild oat EST sequence (GenBank accession: GR359758.1) was identified using the barley and wheat *MLO* gene (Z83834.1 and AF384144.1 respectively) sequences as query sequences in the NCBI BLASTn database. The *A. barbata* sequence was found to share 32% homology to the barley HvMLO protein (GenBank accession: Z83834.1) (Figure S1). As the *MLO* gene family retains high homology across different plant species, primers (Table S1) were designed based on conserved regions between the candidate *A. barbata* sequence (GR359758.1), the barley *HvMLO* (Z83834.1) and the wheat *TaMLO* (AF384144.1) to amplify potential oat *MLO* candidates. A candidate gene was amplified from *A. sativa* cDNA using Phusion High Fidelity Polymerase (New England Biolabs) and primers flanked with Gateway adapter sequences (Table S1). The resulting PCR products were purified using QIA quick PCR Purification Kit (Qiagen), cloned into pDONR207 (Invitrogen) using BP clonase II enzyme mix (Thermo Fisher Scientific) and were verified by sequencing (Macrogen Europe). From this, a partial *AsMLO1* candidate was amplified (Figure S2). With the publication of the *A. sativa* genome, this initial *AsMLO* candidate was identified and labelled *AsMLO1*.

### *in silico* identification of Oat *MLO* family

Protein sequences of previously characterized MLOs were used as a query to identify oat MLOs. To this end, 341 previously characterized MLOs were retrieved from Kusch, *et al.*, (2016) and subsequently queried for presence/absence in the publicly available oat genome (*Avena sativa* – OT3098 v1, PepsiCo, https://wheat.pw.usda.gov/GG3/graingenes_downloads/oat-ot3098-pepsico) using TBLASTN with the following criterion: e-value cut-off <1e-06 and identity>35% (Altschul et al. 1990). From this, 11 *AsMLO* candidate genes were identified (Figure S3). Subsequently, the gene annotation for each of the identified *AsMLO* genes was manually assessed utilizing the publicly available RNAseq data under project number (PRJNA523140) in order to determine the correct exon/intron boundaries for each of these genes. To this end, the PacBio paired-end sequenced reads from the Sequence Read Archive (SRA) were obtained and subsequently trimmed reads to remove adaptors and short reads using FitLong (--min_mean_q 80 --min_length 1000 --keep_percent 90 --target_bases 500000000) (v0.2.0) (github.com/rrwick/Filtlong). Trimmed RNAseq reads were then mapped to the oat genome using minimap2 (default settings) (v.2.17) (Li, 2018). Duplications of the identified *AsMLO* genes were determined using TBLASTN with the following criterion: e-value cut-off <1e-06 and identity>35% (Altschul et al. 1990) (Figure 1).

**Figure 1.**
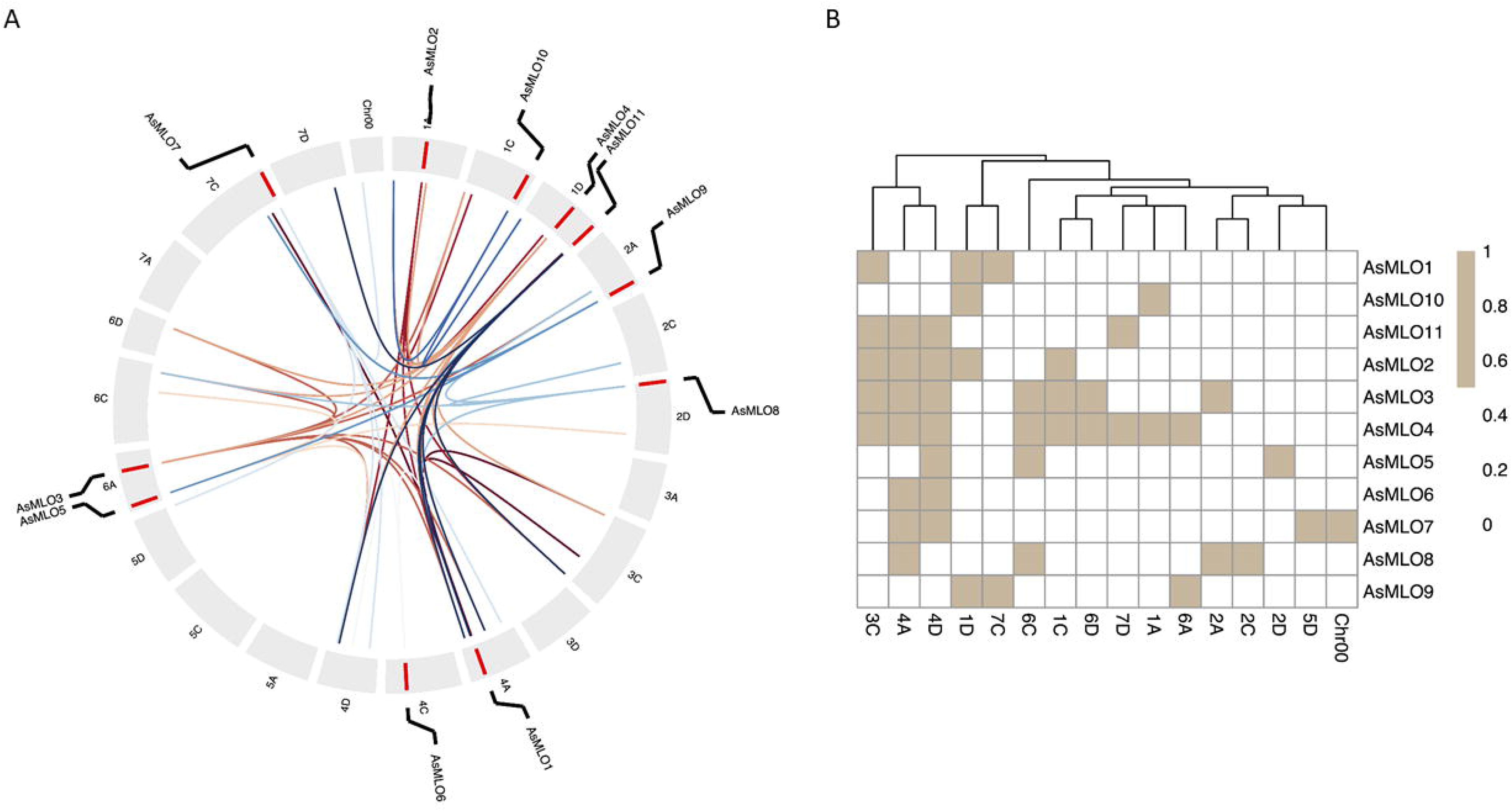
Genomic locations and paralogous *AsMLO* gene pairs in oats. **(A)** Circos diagram showing presence of at least one copy of each *AsMLO* genes across oat chromosomes. The outer lane represents the chromosomes of oat genome. Genomic locations of *AsMLO* genes are depicted as red bars. Paralogous *AsMLO* gene pairs are connected with lines with distinct colours representing *AsMLO* genes. **(B)** Heatmap illustrating presence ‘brown’ or absence ‘white’ of paralogous *AsMLO* pairs across oat genome.

### Multiple Alignment and Comparative Phylogenetic Analysis

The multiple alignment was generated by CLUSTALW using default parameters on the 11 AsMLO proteins and 100 MLO proteins from other plant species, including barley (Hv), wheat (Ta), rice (Os) maize (Zm), *Brachypodium distachyon* (Bd)*, Arabidopsis thaliana* (At)*, Setaria italica* (Si) and tomato (Sl). These protein sequences were obtained from Kusch, *et al.*, (2016). A phylogenetic tree was calculated via the Maximum Likelihood method, using Jones-Taylor-Thornton (JTT) modelling with 500 bootstrap replications in MEGA X (v10.1.7) (Kumar *et al.*, 2018). A second sequence alignment was generated using the plant MLO proteins that clustered into Clade IV. MLO proteins in this clade were previously shown to be involved in powdery mildew susceptibility (Qin *et al.*, 2019). Alignment was generated using CLUSTALW with default parameters and visualized using the ESPript software (http://espript.ibcp.fr/ESPript/ESPript/, Robert and Gouet, 2014).

### Protein Characterisation and Conserved Motifs Analysis

The predicted AsMLO proteins were analysed for physical and chemical characteristics. The number of TM helices were detected using the online server CCTOP (http://cctop.enzim.ttk.mta.hu/?_=) (Dobson, *et al.*, 2015) and TMHMM server (http://www.cbs.dtu.dk/services/TMHMM/) (Möller, *et al.*, 2001), which provides the predicted location of N- and C-terminus of the AsMLOs, either in (cytoplasmic side) or out (extracellular side). The subcellular localization of all AsMLO proteins were determined by the CELLO v2.5 software (http://cello.life.nctu.edu.tw/) (Yu *et al.*, 2006). Domain location and length was determined using the NCBI Conserved Domain Database (https://www.ncbi.nlm.nih.gov/Structure/cdd/wrpsb.cgi) (Lu *et al.*, 2020).

The exon - intron organisations were determined by comparing the coding sequence (CDS) to their corresponding genomic DNA sequences using the online tool GSDS (http://gsds.gao-lab.org/) (Hu *et al.*, 2015). The conserved motifs were detected using the online software MEME v5.3.0 (http://meme-suite.org/tools/meme) (Bailey *et al.*, 2009), with the maximum number of motifs set to 12 and the remaining parameters as system defaults. The results were prepared using TB tool software (Chen *et al.*, 2020).

### qRT-PCR analysis of *AsMLO1* expression in response to powdery mildew

Expression analysis was conducted by real-time quantitative PCR (qRT-PCR) to determine *AsMLO1* response to *Bga* infection. Three-week old oat leaves were inoculated with *Bga*. Two leaves (one leaf from each of two plants) were collected at 0, 8, 12, 24, 48 and 72 hours post-inoculation (hpi) and were flash frozen in liquid nitrogen (N2). Uninfected leaves were also collected and used as controls. RNA from 100 mg of leaf tissue was extracted using the Spectrum Total RNA kit (Sigma-Aldrich). An on-column DNase digestion (Sigma-Aldrich) step was incorporated to remove genomic DNA. Total RNA was quantified using a Nanodrop ND-1000 spectrophotometer. 1 μg of RNA was reverse transcribed using the Thermo Scientific cDNA synthesis kit. Two leaves were pooled within each experiment per time point and three independent experiments were performed (n=6).

qRT-PCR was carried out on the QuantiStudio 7 Flex RT - PCR system (Applied Biosystems) and relative gene expression was calculated as 2-(Ct Target gene – Ct housekeeping gene) using the qRT-PCR threshold cycle (Ct) values. *AsMLO1* gene sequence was used for primer design (Table S1). The oat *28SrRNA* gene was used as a housekeeping control for oats (Jarosová and Kundu, 2010). Data were analysed using the R statistical package (https://www.r-project.org) (R Core Team, 2020) with packages ‘plyr’. ‘multcomp’, ‘emmeans’ and ‘ggplot2’. Analysis of qRT-PCR data was performed using a general linear model. Data were log-transformed prior to analysis. qRT-PCR was performed three times.

## Results

### Identification of *AsMLO* gene family members

To explore all putative *AsMLO* genes in oat, we queried 341 known MLO protein sequences (Kusch, *et al.*, 2016) against the oat genome using TBLASTN and identified 11 putative oat MLO genes. The predicted genes were designated sequentially from *AsMLO1* to *AsMLO11* (Table 1). The gene annotation was manually assessed for all 11 *AsMLO* genes by utilizing publicly available RNA-Seq data under the project number (PRJNA523140) in order to determine the correct exon/intron boundaries for each of these genes. We retrieved the full gene length for each of the *AsMLO* genes except for *AsMLO11* that lacked evidence for the gene model using RNA-Seq support. Thus, *AsMLO11* is referred to as *AsMLO11_partial* in this analysis. *AsMLO* genes were found to be located over nine oat chromosomes (Table 1; Figure 1). *AsMLO3* and *AsMLO5* were found to be located on chromosome 6A; *AsMLO4* and *AsMLO11_partial* located on chromosome 1D; *AsMLO1* on chromosome 4A; *AsMLO2* on chromosome 1A; *AsMLO6* on chromosome 4C; *AsMLO7* on chromosome 7C; *AsMLO8* on chromosome 2D; *AsMLO9* on chromosome 2A; and *AsMLO10* on chromosome 1C. Gene structural analysis, comparing the genomic sequences of the *AsMLO*s to their corresponding coding sequence (CDS) revealed small variations in the number of exons present in each gene, ranging from 11, in *AsMLO11_partial* to 15 exons in *AsMLO2, AsMLO6* and *AsMLO7* (Table 1).

**Table 1.** Features of Identified *AsMLO* genes. *AsMLO* homologs with their respective chromosomal locations, number of exons (determined by the online tool GSDS), protein length (amino acids), domain location and length (determined by NCBI Conserved Domain Database), the number of transmembrane domains (TM; predicted by CCTOP and TMHMM software) and terminus (predicted location of N- and C-terminus, out=extracellular side, in=cytoplasmic side, arrow indicates N-terminus direction to C-terminus), and the subcellular localization of AsMLO proteins (predicted by CELLO 2.5 software).

| MLO name | Chromosome | Exons | Protein length<br>(aa) | MLO domain location | MLO domain length | TM | TM terminus | Subcellular<br>localization |
| --- | --- | --- | --- | --- | --- | --- | --- | --- |
| AsMLO1 | 4A | 13 | 564 | 7 - 501 | 494 | 7 | out → in | Plasma membrane |
| AsMLO2 | 1A | 15 | 562 | 12 - 468 | 456 | 7 | out → in | Plasma membrane |
| AsMLO3 | 6A | 13 | 485 | 13 - 488 | 475 | 7 | out → in | Plasma membrane |
| AsMLO4 | 1D | 14 | 495 | 8 - 478 | 470 | 7 | out → in | Plasma membrane |
| AsMLO5 | 6A | 13 | 505 | 9 - 413 | 404 | 7 | out → in | Plasma membrane |
| AsMLO6 | 4C | 15 | 508 | 5 - 452 | 447 | 7 | out → in | Plasma membrane |
| AsMLO7 | 7C | 15 | 527 | 16 - 459 | 443 | 7 | out → in | Plasma membrane |
| AsMLO8 | 2D | 14 | 491 | 5 - 458 | 453 | 7 | out → in | Plasma membrane |
| AsMLO9 | 2A | 13 | 628 | 5 - 395 | 390 | 7 | out → in | Plasma membrane |
| AsMLO10 | 1C | 13 | 560 | 12 - 514 | 502 | 7 | out → in | Plasma membrane |
| AsMLO11_partial | 1d | 11 | 327 | 1 - 318 | 318 | 4 | out → in | Plasma membrane |

The length of the predicted MLO proteins varies from 327 amino acids (AsMLO11_partial) to 628 amino acids (AsMLO9). Investigation of conserved domains in the protein sequences using a NCBI CDD search resulted in specific hits with the conserved domain model of the MLO superfamily (accession pfam03094) (Devoto *et al.*, 1999), with domain locations ranging from 5 to 514, with the exception of AsMLO11_partial (Table 1).

The subcellular localization predicted by CELLO v2.5 indicates that all AsMLO proteins are localized in the plasma membrane. Seven transmembrane (TM) helices were predicted in AsMLO proteins, with the exception of AsMLO11_partial, which was predicted to have four TM domains due to the partial sequence. The presence of seven TM domains is a typical feature shared among most MLO proteins (Rispail and Rubiales, 2016). Moreover, AsMLO proteins possess an extracellular N-terminus and an cytosolic C-terminus (Table 1).

### *AsMLO* synteny and duplication

Three homeologs of *AsMLO2, AsMLO3, AsMLO4, AsMLO6, AsMLO8* and *AsMLO10* are present on the A, C and D genomes of oats (Figure 1). No homeologs were determined for *AsMLO1, AsMLO5, AsMLO7, AsMLO9* and *AsMLO11_partial*, however duplication of the *AsMLO1* gene is present across the oat chromosomes. Three paralogs of *AsMLO1* are present across the oat chromosomes, which are localized on chromosomes 7C, 1D, and 3C (Figure 1).

Gene duplication serves as an important source of functional divergence, as duplicated genes may gain novel functions or lose original ones contributing to genome evolution (Hughes, 1994; Conant and Wolfe, 2008). The extent of *AsMLO* duplication during the evolution of oats was determined. The most duplicated *AsMLO* gene is *AsMLO4* (9 chromosomes), while the least duplicated one is *AsMLO10* (two chromosomes) (Figure 1). The presence of *AsMLO* gene paralogs may suggest an important role in oat genome evolution.

### Phylogenetic Analysis

Phylogenetic analysis was performed using a total of 100 MLO proteins obtained from Kusch, *et al.*, 2016, and the 11 predicted AsMLO proteins from this study. These 111 proteins clustered into seven phylogenetic clades designated I to VII (Figure 2).

**Figure 2.**
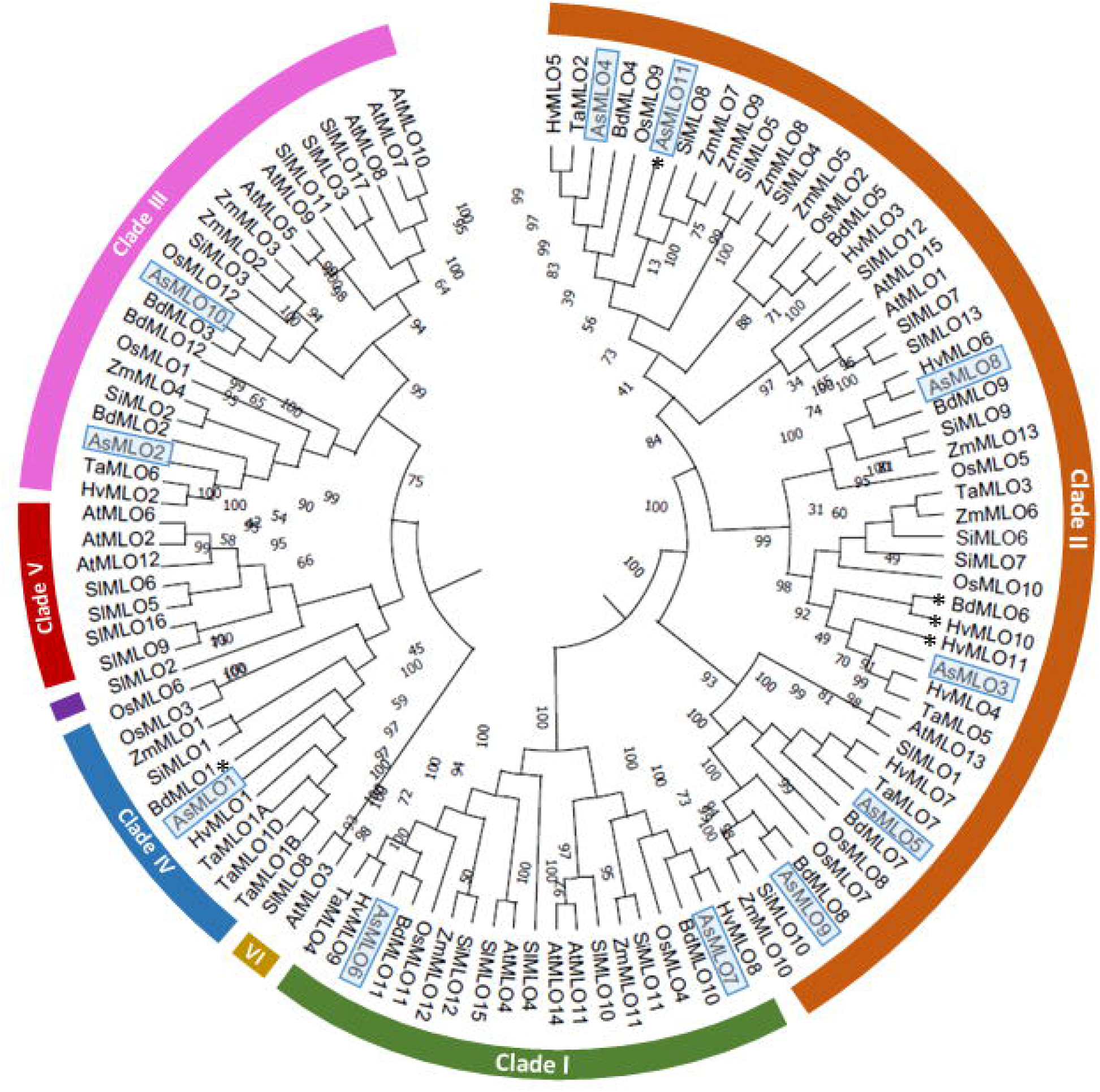
Phylogenetic tree of MLO proteins of oats, *Avena sativa* (As), and other plant species obtained from Kusch, *et al.*, (2016). Plant species include *Arabidopsis thaliana* (At), *Brachypodium distachyon* (Bd), *Hordeum vulgare*, barley *(Hv*), *Oryza sativa*, rice (Os), *Setaria italica* (Si), *Solanum lycopersicum*, tomato (Sl), *Triticum aestivum*, wheat (Ta) and *Zea mays*, maize (Zm). The phylogenetic tree was calculated *via* the Maximum Likelihood method, using JTT modelling with 500 bootstrap replications in MEGA X. MLO proteins were grouped into 7 different clades, according to Kusch, *et al.*, 2016. The percentage of trees in which the associated taxa clustered together is shown next to the branches. Purple marker indicates Clade VII. Blue boxes highlight AsMLOs. Proteins with asterisks indicate; BdMLO1 and BdMLO6 are isoform 1; AsMLO11, HvMLO10 and HvMLO11 are partial sequences.

The AsMLO proteins were found within Clades I to IV; Clade I contained AsMLO6 and AsMLO7; Clade II contained AsMLO3, AsMLO4, AsMLO5, AsMLO8, AsMLO9 and AsMLO11_partial; Clade III contained AsMLO2 and AsMLO10; and Clade IV contains AsMLO1. Notably, AsMLO1 clustered within Clade IV together with the BdMLO1_isoform1 from *Brachypodium distachyon* and the *Hordeum vulgare* (barley) HvMLO1. HvMLO is associated with powdery mildew susceptibility (Piffanelli *et al.*, 2002). However, expression analysis of *BdMLO1*_isoform1 in response to powdery mildew infection has not been demonstrated (Ablazov and Tombuloglu, 2016).

Multiple sequence alignment of the Clade IV MLO proteins was performed to determine the common structural features between potential powdery mildew susceptibility factors. AsMLO1 and the MLO homologs belonging to Clade IV share highly conserved TM domains. In addition, the CaMBD domain and peptide domains I and II within the C-terminus of MLO proteins are conserved (Figure 3) (Kim *et al.*, 2002; Panstruga, 2005).

**Figure 3.**
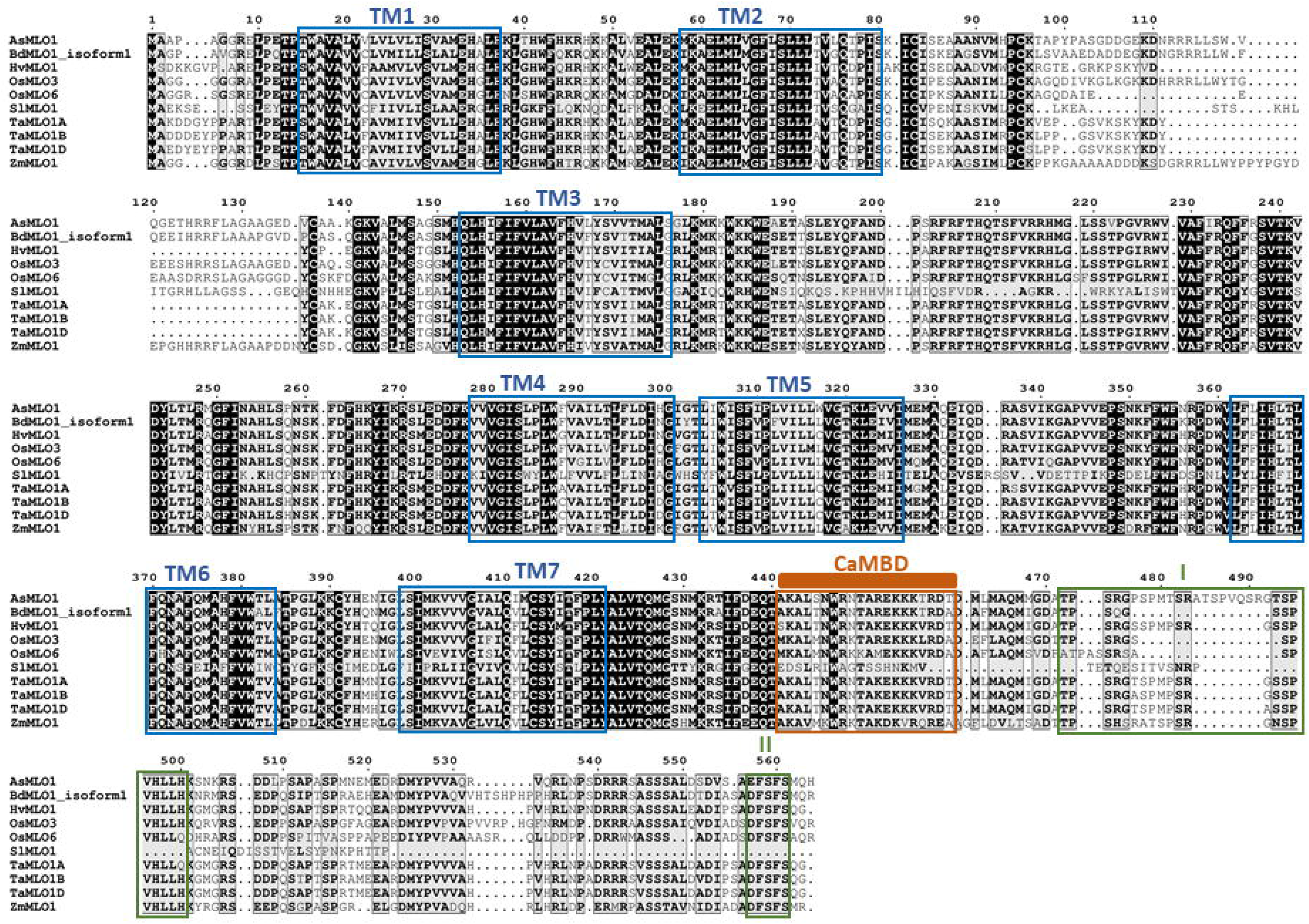
Sequence alignment of MLO proteins in Clade IV; MLOs previously shown to be involved in powdery mildew susceptibility. Black shading and its intensity indicate conserved sites and the degree of conservation. The alignment was generated by CLUSTALW using default parameters. Blue boxes indicate the positions of the seven TM domains (TM1-7) adapted from Kusch, *et al.*, 2016; orange box indicates the approximate position of the calmodulin-binding domain (CaMBD) inferred from Kim, *et al.*, (2002); green boxes indicate the peptide domains I and II, inferred from Panstruga, (2005) and Chen, *et al.*, (2020).

### Conserved Amino Acids and Motif Analysis of AsMLO Proteins

The conserved motif analysis completed through the MEME tool generated 14 motifs with lengths varying between 6 and 50 amino acids. Motifs 1, 2, 3, 4, 8 and 9 were found to be present in all AsMLO proteins (Table 2, Figure 4). The amino acids of these motifs were compared to the conserved motifs from the land plant MLO analysis completed by Kusch, *et al.*, (2016). Motif 1 contains a derivative of the highly conserved sub - motif [FY]DF which corresponds to the Cytoplasmic loop 2 domain and possible carbohydrate metabolism. Motif 2 contains a FWF motif that corresponds to Cytoplasmic loop 3 and ion transport. Motifs 3 and 4 contain derivatives of the WxxWE and SFFKQF with TLRxGFI motifs, respectively. These motifs correspond to Cytoplasmic loop 2 but have no suggested function. Motif 6 contains a derivative of the EALEK motif, which corresponds to Cytoplasmic loop 1 and fatty acid metabolism. While Motif 7 contains TPTW, associated with TM domain 1. It is likely both Motif 6 and 7 are present in all AsMLO proteins but are missing in AsMLO11_partial due to the partial sequence.

**Table 2.**
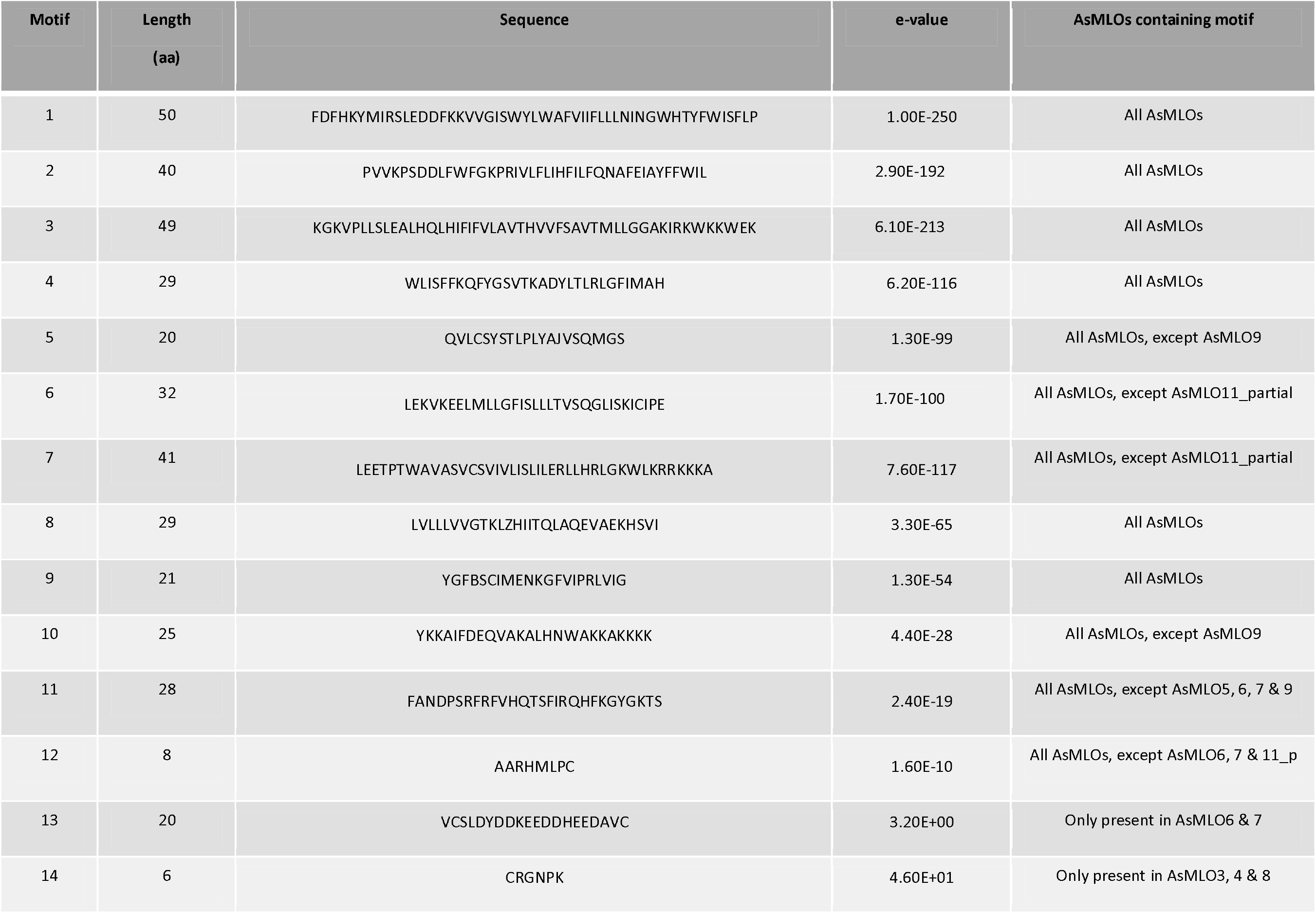
Conserved motifs in predicted AsMLO proteins.

**Figure 4.**
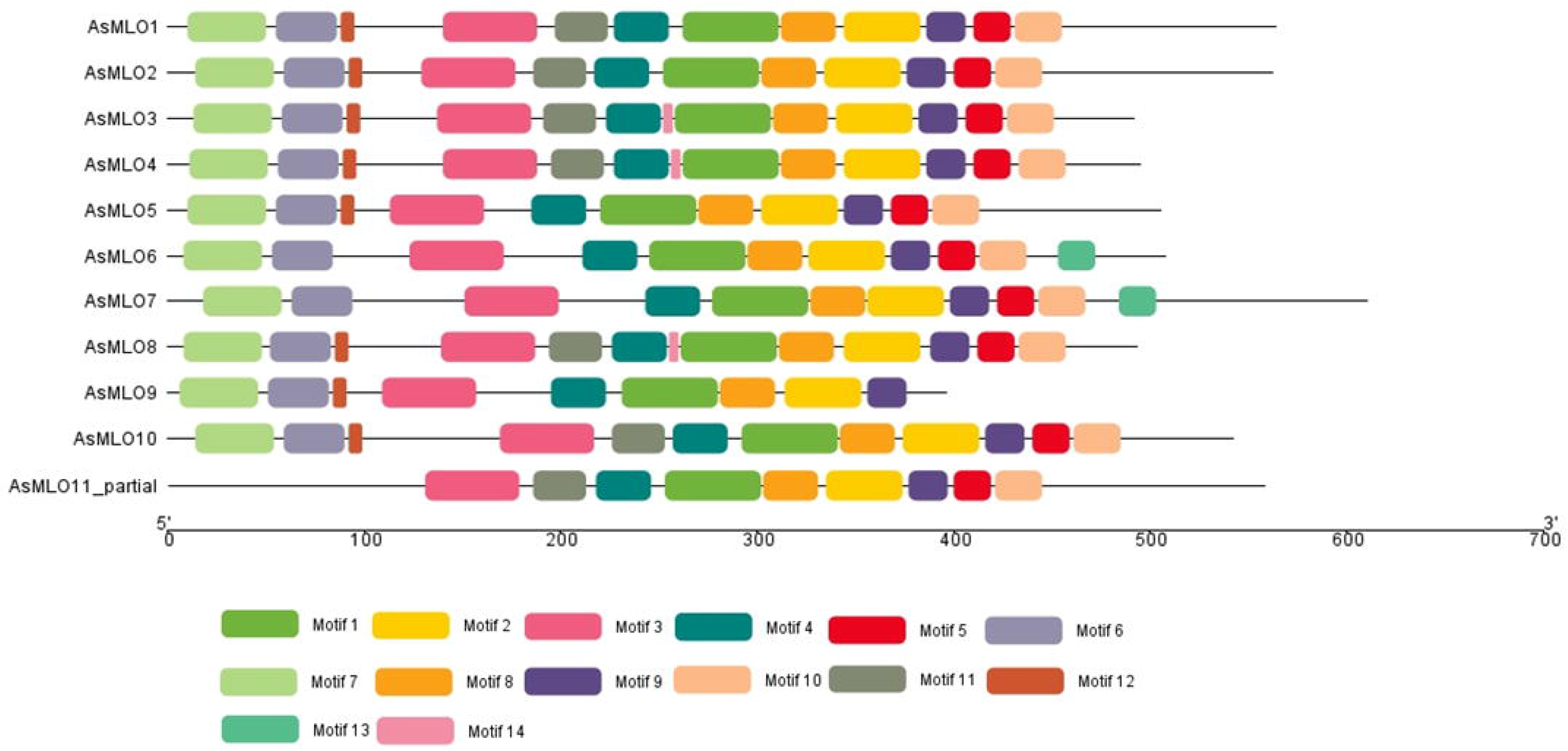
Schematic distribution of conserved motifs within the AsMLO proteins. The conserved motifs were detected using the online software MEME v5.3.0 (http://meme-suite.org/tools/meme) (Bailey *et al.*, 2009), with the maximum number of motifs set to 14 and the remaining parameters as system defaults. The results were prepared using TB tool software (Chen *et al.*, 2020). Each coloured box represents an individual motif sequence, outlined in Table 2.

### Expression Analysis of *AsMLO1* in response to *Bga* infection

*Bga* infection was microscopically examined at the time points 0, 8, 12, 24, 48 and 72 hours post infection (hpi) to visualize each infection stage of powdery mildew in oats. The microscopy demonstrates that *Bga* penetration occurs between 8 and 12 hpi (Figure 5A). To determine a potential response of *AsMLO1* to powdery mildew infection, we examined expression at these time points of powdery mildew infection using qRT - PCR (Figure 5B).

**Figure 5.**
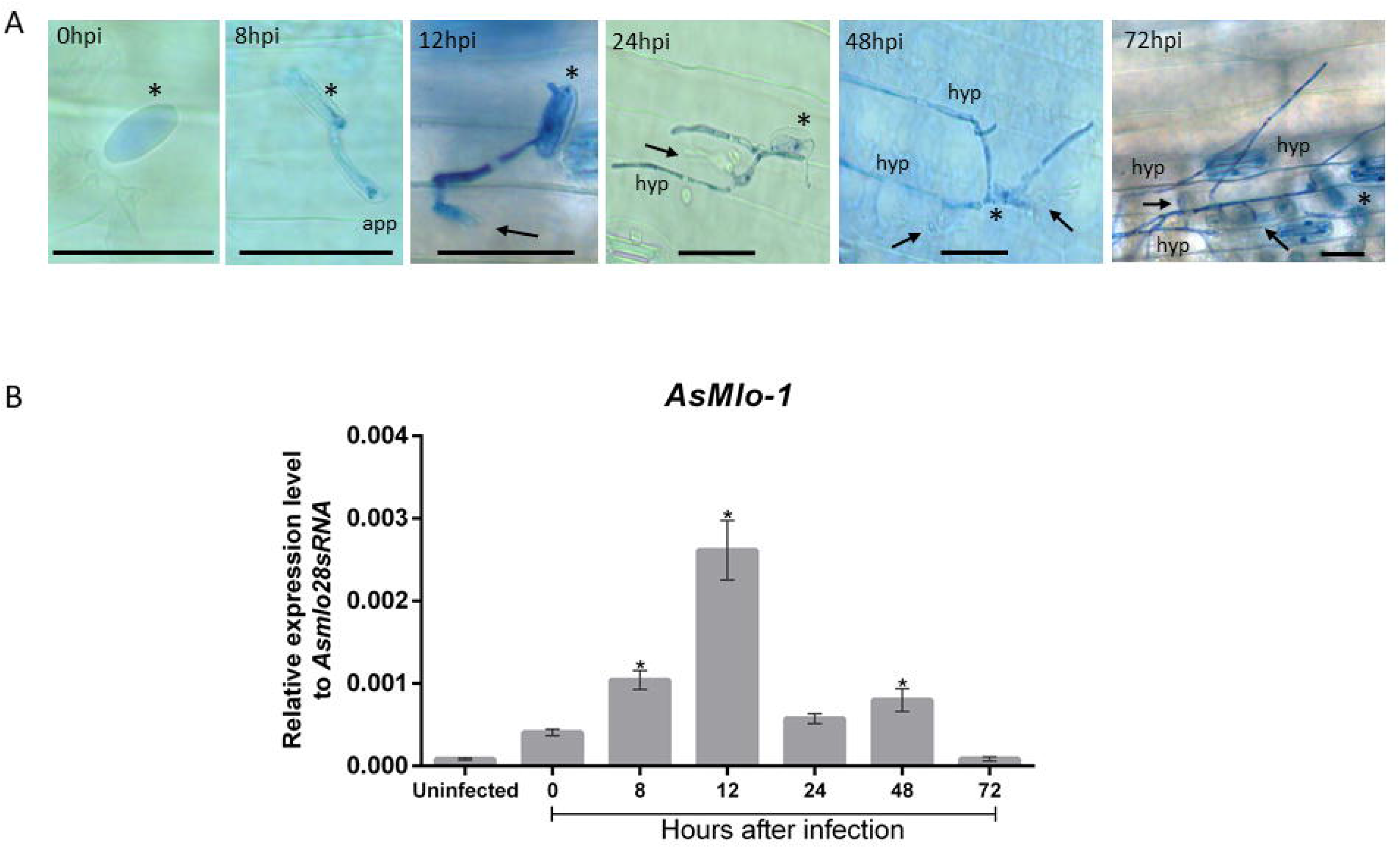
**(A)** *Blumeria graminis* f. spp. *avenae, (Bga)* progression on oat leaves, visualized with trypan blue at 0, 8, 12, 24, 48 and 72 hpi. Scale black bars = 50μm. Black asterisks = conidium; app = appressorium; black arrow = haustoria, hyp = hyphae. **(B)** Gene expression analysis of *AsMLO1* at 0, 8, 12, 24, 48 and 72 hpi of oat (cultivar Barra) with *Bga.* RNA from two leaves (one leaf from each of two plants) were collected at uninfected 0, 8, 12, 24, 48 and 72 hpi was extracted and quantified using a Nanodrop ND-1000 spectrophotometer. 1 μg of RNA was reverse transcribed using the Thermo Scientific cDNA synthesis kit. Two leaves were pooled within each experiment per time point and three independent experiments were performed (n=6). cDNA from all three experimental replicates were used on each 96 – well qRT – PCR plate with specific primers for the candidate gene. The expressions levels of *As*28S*rRNA* were used to normalize expression levels of *AsMLO1.* Three independent qRT – PCR plates were performed. The bars represent the mean relative expression ± SEM. The asterisk on top of the bar represents significant differences determined by ANOVA(\**P*<0.05).

A significant increase in *AsMLO1* gene expression was observed in oat leaf tissue at 8 hours post *Bga* infection and 12 and 48 hpi, when expression is compared with uninfected leaves. Relative gene expression peaked at 12 hours, and then decreased at 24 hours. Transcript levels significantly increased again at 48 hours and finally decreased by 72 hours (Figure 5B).

## Discussion

Powdery mildew is one of the major fungal diseases that affects oat production globally (Sánchez-Martín *et al.*, 2016). Mutations in the barley *MLO* gene were found to be key in providing durable genetic resistance to powdery mildew in barley (Jørgensen, 1992). The *MLO* gene family has become an important focus for disease resistance research due to the gene family’s presence in all higher plants and the role specific gene members play as susceptibility factors for powdery mildew disease (Polanco *et al.*, 2018). In this study, we report the identification of the *AsMLO* family from the oat genome through an *in silico* approach. Eleven *AsMLO* members were found which correspond with the number of *MLO*s in other monocot species. For example, 11 *MLO*s were found in barley and *B. distachyon* while, 12 *MLO*s were found in rice (Ablazov and Tombuloglu, 2016; Kusch, *et al.*, 2016). Homeologs of *AsMLO2, AsMLO3, AsMLO4, AsMLO6, AsMLO8* and *AsMLO10* are present on the A, C and D genomes of oats (Figure 1) which is a hexaploid. However, homeologs were not found for *AsMLO1, AsMLO5, AsMLO7, AsMLO9* and *AsMLO11_partial*. Similar to our findings, the octoploid strawberry (*Fragaria*×*ananassa*) had varying numbers of homeologs for each of the 20 *FaMLOs* identified ranging from none through to seven homeologs (Tapia *et al*., 2020). Previously, the homeologs TaMlo-A1, TaMlo-B1 and TaMlo-D1 were identified in wheat which are associated with powdery mildew susceptibility (Elliot *et al.*, 2002).

The deduced proteins of the AsMLOs were also found to shared key attributes with MLO proteins in other plant species (Table 1). These attributes include the presence of seven predicted TM domains and localisation in the plasma membrane. Some studies have suggested that the seven TM domains can vary from 2 to 10 (Deshmukh, *et al.*, 2014; Chen *et al.*, 2020). However, with respect to the present study, AsMLO proteins were predicted to have seven, with an extracellular N-terminus and a cytoplasmic C-terminus. The only exception to this was AsMLO11_partial, but this is because a full sequence was not obtained.

The multiple sequence alignment of Clade IV proteins showed that AsMLO1 shared seven highly conserved TM domains, a predicted CaMBD domain (Kim *et al.*, 2002) and peptide domains I and II at the C-terminus (Panstruga, 2005) (Figure 3). The CaMBD likely supports MLO-mediated susceptibility by recognising the changes of Ca+ ions during fungal penetration (Chen *et al.*, 2020). Two peptide domains (I and II were also detected in AsMLO1, downstream of the CaMBD that may have a function in modulating powdery mildew susceptibility (Panstruga, 2005). These domains in AsMLO1 (TPSRGPSPMTSRATSPVQSRGTSPVHLLH (I) and EFSFS (II)) showed high conservation to Clade IV proteins (Figure 3) but were not detected as shared motifs with any other AsMLO protein (Table 2). This evidence supports AsMLO1 as a powdery mildew associated MLO in oats.

Our phylogenetic analyses of AsMLO proteins with different monocot and dicot plant species produced seven clades, of which AsMLOs were represented within four clades (Clade I to IV). This conforms with other studies that conclude MLO proteins from monocot plant species are found in clades I - IV only (Kusch, *et al.*, 2016). All monocot MLOs that have been functionally verified as susceptibility factors have been shown to cluster within Clade IV (Qin *et al.*, 2019). Previous reports stated *B. distachyon* ecotypes expressed high levels of resistance to strains of *B. graminis* (Fitzgerald *et al.*, 2015). This is potentially due to a low copy of MLO paralogs in *B. distachyon* (Ablazov and Tombuloglu, 2016). However, *BdMLO* has only been demonstrated to be associated with *F. graminearium* infection to date (Ablazov and Tombuloglu, 2016).

The presence of AsMLO1 in Clade IV provides an indication of its possible function as a powdery mildew susceptibility factor in oats. For these reasons, AsMLO1 was selected for expression analysis. The other clades that contain AsMLOs proteins (I - III) have been shown to include MLOs that are expressed in other plant tissue and perhaps involved in other physiological functions (Figure 2) (Feechan *et al*., 2009; Liu and Zhu, 2008; Konishi, *et al.*, 2010; Acevedo-Garcia, *et al.*, 2014; Kusch, *et al.*, 2016).

The main characteristic of powdery mildew associated MLOs is the up-regulation of their expression once challenged with *B. graminis* (Feechan *et al*., 2008; Iovieno *et al.*, 2015). Previous research on *HvMLO1* expression on *Bgh* infected barley determined peak expression by 6 hpi; a rapid up-regulation that correlated with *Bgh* attempted penetration (Piffanelli *et al.*, 2002). We were able to demonstrate significant upregulation of *AsMLO1* expression at 8 hpi in response to *Bga* challenge and expression peaked at the later time point of 12 hpi (Figure 5). This is in line with *Bga* penetration of leaf cells prior to the formation of a mature feeding structure (haustoria) and hyphal elongation.

In this study, we identified the *AsMLO* family and AsMLO1, which shares similarities to other *MLO* genes associated with powdery mildew susceptibility, in regard to sequence homology and protein characteristics. The findings reported in this study provide knowledge that can be used to generate *mlo* oat varieties in the future that are resistant to powdery mildew.

## Acknowledgements

This research was supported by a Teagasc Walsh Scholarship.

## Author Contributions

A.R, A.F, S.K and J.F designed experiments. H.G performed genome analysis. A.R and S.J.K performed initial primer design and cloning. A.R analysed the data. S.J.K performed qRT-PCR and statistical analysis. A.T performed cDNA synthesis. A.R, A.F, H.G and S.K wrote the manuscript with input from S.J.K and A.T.

## Supporting Information

**Figure S1.** Alignment of *Avena barbata MLO* candidate, barley (HvMLO) and wheat MLO (TaMLO) protein sequences. Alignment was performed using Multalin (v5.4.1) and visualized using ESPrint software. Genbank accession numbers of sequences used for the alignment: *A. barbata* (GR359758.1) translated using ExPASy; barley (CAB06083) and wheat (AAK60566).

**Figure S2:** Nucleotide alignment of the partial *AsMLO* candidate (AsMLO1_c) that was cloned using primers based on the *Avena barbata, HvMLO* and *TaMLO* sequences. Alignment was performed using Multalin (v5.4.1) and visualized using ESPrint software. Genbank accession numbers of *MLO* nucleotide sequences used in alignment: *A. barbata* (GR359758.1); *HvMLO* (Z83834.1); and *TaMLO* (AF384144.1).

**Figure S3.** Alignment of 11 AsMLO protein sequences. Alignment was performed using Multalin (v5.4.1).

## References

Ablazov, A. and Tombuloglu, H. (2016) ‘Genome-wide identification of the mildew resistance locus O (MLO) gene family in novel cereal model species Brachypodium distachyon’, European Journal of Plant Pathology, 145(2), pp. 239–253. doi: 10.1007/s10658-015-0833-2.

Acevedo-Garcia, J., Spencer, D., Thieron, H., Reinstädler, A., Hammond-Kosack, K., Phillips, A. L. and Panstruga, R. (2017) ‘mlo-based powdery mildew resistance in hexaploid bread wheat generated by a non-transgenic TILLING approach’, Plant Biotechnology Journal, 15(3), pp. 367–378. doi: https://doi.org/10.1111/pbi.12631.

Altschul, S. F., Gish, W., Miller, W., Myers, E. W., Lipman, D. J. (1990) ‘Basic local alignment search tool’, Journal of Molecular Biology, 215(3), pp. 403–410.

Appiano, M., Catalano, D., Santillán Martínez, M., Lotti, C., Zheng, Z., Visser, R. G. F., Ricciardi, L., Bai, Y. and Pavan, S. (2015) ‘Monocot and dicot MLO powdery mildew susceptibility factors are functionally conserved in spite of the evolution of class-specific molecular features’, BMC Plant Biology, 15(1), p. 257. doi: 10.1186/s12870-015-0639-6.

Bailey, T. L., Boden, M., Buske, F. A., Frith, M., Grant, C. E., Clementi, L., Ren, J., Li, W. W. and Noble, W. S. (2009) ‘MEME SUITE: tools for motif discovery and searching.’, Nucleic acids research, 37(Web Server issue), pp. W202–8. doi: 10.1093/nar/gkp335.

Chen, C., Chen, H., Zhang, Y., Thomas, H.R., Frank, M. H., He, Y. and Xia, R. (2020) ‘TBtools - an integrative toolkit developed for interactive analyses of big biological data’, bioRxiv, p. 289660. doi: 10.1101/289660.

Chen, L., Chen, C., Liu, J., Liu, Z., Xia, P., Yuan, X. and Ning, Y. (2020) ‘Genome-wide identification and expression analysis of the MLO gene family reveal a candidate gene associated with powdery mildew susceptibility in bitter gourd (Momordica charantia)’, European Journal of Plant Pathology. doi: 10.1007/s10658-020-02152-0.

Clifford, B. C. (1995) ‘Diseases, pests and disorders of oats’, The Oat Crop: Production and Utilization, pp. 252–278. doi: 10.1007/978-94-011-0015-1_9.

Cui, F., Wu, H., Safronov, O., Zhang, P., Kumar, R., Kollist, H., Salojärvi, Ja., Panstruga, R. and Overmyer, K. (2018) ‘Arabidopsis MLO2 is a negative regulator of sensitivity to extracellular reactive oxygen species’, Plant, Cell & Environment, 41(4), pp. 782–796. doi: https://doi.org/10.1111/pce.13144.

Das, A., Sharma, N. and Prasad, M. (2019) ‘CRISPR/Cas9: A Novel Weapon in the Arsenal to Combat Plant Diseases’, Frontiers in Plant Science, p.2008. Available at: https://www.frontiersin.org/article/10.3389/fpls.2018.02008.

Deshmukh, R., Singh, V. K. and Singh, B. D. (2014) ‘Comparative phylogenetic analysis of genome-wide Mlo gene family members from Glycine max and Arabidopsis thaliana’, Molecular Genetics and Genomics, 289(3), pp. 345–359. doi: 10.1007/s00438-014-0811-y.

Dobson, L., Reményi, I. and Tusnády, G. E. (2015) ‘CCTOP: a Consensus Constrained TOPology prediction web server’, Nucleic acids research, 43(W1), pp. W408–W412. doi: 10.1093/nar/gkv451.

Elliott, C., Zhou, F., Spielmeyer, W., Panstruga, R. and Schulze-Lefert, P. (2002) ‘Functional conservation of wheat and rice Mlo orthologs in defense modulation to the powdery mildew fungus.’, Molecular plant-microbe interactions◻: MPMI, 15(10), pp. 1069–1077. doi: 10.1094/MPMI.2002.15.10.1069.

Feechan, A., Jermakow, A. M., Torregrosa, L., Panstruga, R. and Dry, I. B. (2008) ‘Identification of grapevine MLO gene candidates involved in susceptibility to powdery mildew’, Functional Plant Biology, 35(12), pp. 1255–1266. Available at: https://doi.org/10.1071/FP08173.

Fitzgerald, T. L., Powell, J., Schneebeli K., Hsia, M. M., Gardiner, D. M., Bragg, J. N., McIntyre, C. L., Manners, J. M., Ayliffe, M., Watt, M., Vogel, J. P., Henry, R. and Kazan, K. (2015) ‘Brachypodium as an emerging model for cereal-pathogen interactions’, Annals of botany, 115(5), pp. 717–731. doi:10.1093/aob/mcv010.

Hu, B., Jin, J., Guo, A.-Y., Zhang, H., Luo, J., Gao, G. (2015) ‘GSDS 2.0: an upgraded gene feature visualization server’, Bioinformatics, 31(8), pp. 1296–1297. doi: 10.1093/bioinformatics/btu817.

Iovieno, P., Andolfo, G., Schiavulli, A., Catalano, D., Ricciardi, L., Frusciante, L., Ercolano, M. R. and Pavan, S. (2015) ‘Structure, evolution and functional inference on the Mildew Locus O (MLO) gene family in three cultivated Cucurbitaceae spp.’, BMC Genomics, 16(1), p. 1112. doi: 10.1186/s12864-015-2325-3.

Isidro-Sánchez, J., Prats, E., Howarth, C., Langdon, T. and Montilla-Bascón, G. (2020) ‘Genomic Approaches for Climate Resilience Breeding in Oats’, in Genomic Designing of Climate-Smart Cereal Crops, pp. 133–169. doi: 10.1007/978-3-319-93381-8_4.

Jarosová, J. and Kundu, J. K. (2010) ‘Validation of reference genes as internal control for studying viral infections in cereals by quantitative real-time RT-PCR’, BMC Plant Biology., 10, p. 146. doi: 10.1186/1471-2229-10-146.

Jørgensen, I. H. (1992) ‘Discovery, characterization and exploitation of Mlo powdery mildew resistance in barley’, Euphytica, 63(1), pp. 141–152. doi: 10.1007/BF00023919.

Kim, M. C., Lee, S. H., Kim, J. K., Chun, H. J., Choi, M. S., Chung, W. S., Moon, B. C, Kang, C. H., Park, C. Y., Yoo, J. H., Kang, Y. H., Koo, S. C., Koo, Y. D., Jung, J. C., Kim, S. T., Schulze-Lefert, P., Lee, S. Y. and Cho M. J. (2002) ‘Mlo, a modulator of plant defense and cell death, is a novel calmodulin-binding protein. Isolation and characterization of a rice Mlo homologue.’, The Journal of Biological Chemistry, 277(22), pp. 19304–19314. doi:10.1074/jbc.M108478200.

Koch, E. and Slusarenko, A. (1990) ‘Arabidopsis Is Susceptible to Infection by a Downy Mildew Fungus’, The Plant Cell, 2, pp. 437–445. doi: 10.1105/tpc.2.5.437.

Konishi, S., Sasakuma, T. and Sasanuma, T. (2010) ‘Identification of novel *Mlo* family members in wheat and their genetic characterization’, Genes & Genetic Systems, 85(3), pp. 167–175. doi: 10.1266/ggs.85.167.

Kumar, S., Stecher, G., Li, M., Knyaz, C. and Tamura, K. (2018) ‘MEGA X: Molecular Evolutionary Genetics Analysis across Computing Platforms.’, Molecular biology and evolution, 35(6), pp. 1547–1549. doi: 10.1093/molbev/msy096.

Kusch, S. and Panstruga, R. (2017) ‘mlo-Based Resistance: An Apparently Universal “Weapon” to Defeat Powdery Mildew Disease’, Molecular Plant-Microbe Interactions, 30(3), pp. 179–189. doi: 10.1094/MPMI-12-16-0255-CR.

Kusch, S., Pesch, L. and Panstruga, R. (2016) ‘Comprehensive Phylogenetic Analysis Sheds Light on the Diversity and Origin of the MLO Family of Integral Membrane Proteins’, Genome Biology and Evolution, 8(3), pp. 878–895. doi: 10.1093/gbe/evw036.

Li, H. (2018) ‘Minimap2: pairwise alignment for nucleotide sequences’, Bioinformatics, 34(18), pp.3094–3100

Liu, Q. and Zhu, H. (2008) ‘Molecular evolution of the MLO gene family in Oryza sativa and their functional divergence’, Gene, 409(1), pp. 1–10. doi: https://doi.org/10.1016/j.gene.2007.10.031.

Lu, S. et al. (2020) ‘CDD/SPARCLE: the conserved domain database in 2020.’, Nucleic acids research, 48(D1), pp. D265–D268. doi: 10.1093/nar/gkz991.

McGrann, G. R. D., Stavrinides, A., Russell, J., Corbitt, M. M., Booth, A., Chartrain, L., Thomas, W. T. B. and Brown, J. K. M. (2014) ‘A trade off between mlo resistance to powdery mildew and increased susceptibility of barley to a newly important disease, Ramularia leaf spot.’, Journal of experimental botany, 65(4), pp. 1025–1037. doi: 10.1093/jxb/ert452.

Möller, S., Croning, M. D. and Apweiler, R. (2001) ‘Evaluation of methods for the prediction of membrane spanning regions.’, Bioinformatics (Oxford, England), 17(7), pp. 646–653. doi: 10.1093/bioinformatics/17.7.646.

Nguyen, V. N. T., Vo, K. T. X., Park, H., Jeon, J.-S. and Jung, K.-H. (2016) ‘A Systematic View of the MLO Family in Rice Suggests Their Novel Roles in Morphological Development, Diurnal Responses, the Light-Signaling Pathway, and Various Stress Responses’, Frontiers in Plant Science, p. 1413. Available at: https://www.frontiersin.org/article/10.3389/fpls.2016.01413.

Niks, R. E. and Rubiales, D. (2002) ‘Potentially durable resistance mechanisms in plants to specialised fungal pathogens’, Euphytica, 124(2), pp. 201–216. doi: 10.1023/A:1015634617334.

Ociepa, T., Okoń, S., Nucia, A., Leśniowska-Nowak, J., Paczos-Grzęda, E. and Bisaga, M. (2020) ‘Molecular identification and chromosomal localization of new powdery mildew resistance gene Pm11 in oat’, Theoretical and Applied Genetics, 133(1), pp. 179–185. doi: 10.1007/s00122-019-03449-3.

Okoń, S. M. (2015) ‘Effectiveness of resistance genes to powdery mildew in oat’, Crop Protection, 74, pp. 48–50. doi: 10.1016/J.CROPRO.2015.04.004.

Panstruga, R. (2005) ‘Discovery of Novel Conserved Peptide Domains by Ortholog Comparison within Plant Multi-Protein Families’, Plant Molecular Biology, 59(3), pp. 485–500. doi: 10.1007/s11103-005-0353-0.

Panstruga, R. and Kuhn, H. (2019) ‘Mutual interplay between phytopathogenic powdery mildew fungi and other microorganisms’, Molecular Plant Pathology, 20(4), pp. 463–470. doi: 10.1111/mpp.12771.

Pavan, S., Jacobsen, E., Visser, R. G. F., Bai, Y., (2010) ‘Loss of susceptibility as a novel breeding strategy for durable and broad-spectrum resistance’, Molecular breeding◻: new strategies in plant improvement, 25(1), pp. 1–12. doi: 10.1007/s11032-009-9323-6.

PepsiCo (2020) Avena sativa – OT3098 v1, GrainGenes - A Database for Triticeae and Avena. Available at: https://wheat.pw.usda.gov/GG3/graingenes_downloads/oat-ot3098-pepsico.

Pessina, S., Lenzi, L., Perazzolli, M., Campa, M., Dalla Costa, L., Urso, S., Valè, G., Salamini, F., Velasco, R. and Malnoy, M. (2016) ‘Knockdown of MLO genes reduces susceptibility to powdery mildew in grapevine’, Horticulture Research, 3(1), p. 16016. doi: 10.1038/hortres.2016.16.

Piffanelli, P., Zhou, F., Casais, C., Orme, J., Jarosch, B., Schaffrath, U., Collins, N. C., Panstruga, R. and Schulze-Lefert, P. (2002) ‘The barley MLO modulator of defense and cell death is responsive to biotic and abiotic stress stimuli’, Plant physiology, 129(3), pp. 1076–1085. doi: 10.1104/pp.010954.

Polanco, C., Sáenz de Miera, L. E., Bett, K. and Pérez de la Vega, M. (2018) ‘A genome-wide identification and comparative analysis of the lentil MLO genes’, PLOS ONE, 13(3), p. e0194945. Available at: https://doi.org/10.1371/journal.pone.0194945.

Qin, B. et al. (2019) ‘Identification and Characterization of a Potential Candidate Mlo Gene Conferring Susceptibility to Powdery Mildew in Rubber Tree’, Phytopathology, 109(7), pp. 1236–1245. doi: 10.1094/PHYTO-05-18-0171-R.

R Core Team (2020) ‘R: A Language and Environment for Statistical Computing’. Vienna, Austria: R Foundation for Statistical Computing. Available at: https://www.r-project.org/.

Rispail, N. and Rubiales, D. (2016) ‘Genome-wide identification and comparison of legume MLO gene family’, Scientific reports, 6, p. 32673. doi: 10.1038/srep32673.

Robert, X. and Gouet, P. (2014) ‘Deciphering key features in protein structures with the new ENDscript server’, Nucleic Acids Research, 42(W1), pp. W320–W324. doi: 10.1093/nar/gku316.

Roderick, H. W., Jones, E. R. L. and Šebesta, J. (2000) ‘Resistance to oat powdery mildew in Britain and Europe: a review’, Annals of Applied Biology, 136(1), pp. 85–91. doi: 10.1111/j.1744-7348.2000.tb00012.x.

Sánchez-Martín, J., Montilla-Bascón, G., Mur, L. A. J., Rubiales, D. and Prats, E. (2016) ‘Compromised Photosynthetic Electron Flow and H2O2 Generation Correlate with Genotype-Specific Stomatal Dysfunctions during Resistance against Powdery Mildew in Oats’, Frontiers in Plant Science, p. 1660. Available at: https://www.frontiersin.org/article/10.3389/fpls.2016.01660.

Tapia R. R., Barbey C. R., Chandra S., Folta, K. M., Whitaker, V. M. and Lee, S.(2020) ‘Genome-Wide Identification and Characterization of *MLO* Gene Family in Octoploid Strawberry (Fragaria ×ananassa)’ bioRxiv, p.p 2020.02.03.932764 DOI: 10.1101/2020.02.03.932764.

Takamatsu, S. and Matsuda, S. (2004) ‘Estimation of molecular clocks for ITS and 28S rDNA in Erysiphales’, Mycoscience, 45(5), pp. 340–344. doi:https://doi.org/10.1007/S10267-004-0187-7.

Twamley, T., Gaffney, M. and Feechan, A. (2019) ‘A Microbial Fermentation Mixture Primes for Resistance Against Powdery Mildew in Wheat’, Frontiers in Plant Science, p.1241. Available at: https://www.frontiersin.org/article/10.3389/fpls.2019.01241.

Underwood, W. and Somerville, S. C. (2008) ‘Focal accumulation of defences at sites of fungal pathogen attack’, Journal of Experimental Botany, 59(13), pp. 3501–3508. doi: 10.1093/jxb/ern205.

Várallyay, É., Giczey, G. and Burgyán, J. (2012) ‘Virus-induced gene silencing of Mlo genes induces powdery mildew resistance in Triticum aestivum’, Archives of Virology, 157(7), pp. 1345–1350. doi: 10.1007/s00705-012-1286-y.

Wan, D.-Y., Guo, Y., Cheng, Y., Hu, Y., Xiao, S., Wang, Y. and Wen, Y.-Q. (2020) ‘CRISPR/Cas9-mediated mutagenesis of VvMLO3 results in enhanced resistance to powdery mildew in grapevine (Vitis vinifera)’, Horticulture Research, 7(1), p. 116. doi: 10.1038/s41438-020-0339-8.

Wang, Y., Cheng, X., Shan, Q., Zhang, Y., Liu, J., Gao, C. and Qiu, J.-L. (2014) ‘Simultaneous editing of three homoeoalleles in hexaploid bread wheat confers heritable resistance to powdery mildew’, Nature Biotechnology, 32(9), pp. 947–951. doi: 10.1038/nbt.2969.

Yu, C.-S., Chen, Y. C., Lu, C. H. and Hwang, J. K. (2006) ‘Prediction of protein subcellular localization.’, Proteins, 64(3), pp. 643–651. doi: 10.1002/prot.21018.

Yu, G., Wang, X., Chen, Q., Cui, N., Yu, Y. and Fan, H. (2019) ‘Cucumber Mildew Resistance Locus O Interacts with Calmodulin and Regulates Plant Cell Death Associated with Plant Immunity’, International Journal of Molecular Sciences, 20(12), p. 2995. doi: 10.3390/ijms20122995.

Yu, J. and Herrmann, M. (2006) ‘Inheritance and mapping of a powdery mildew resistance gene introgressed from Avena macrostachya in cultivated oat’, Theoretical and Applied Genetics, 113(3), pp. 429–437. doi: 10.1007/s00122-006-0308-0.

Zimny, T., Sowa, S., Tyczewska, A. and Twardowski, T. (2019) ‘Certain new plant breeding techniques and their marketability in the context of EU GMO legislation – recent developments’, New Biotechnology, 51, pp. 49–56. doi: https://doi.org/10.1016/j.nbt.2019.02.003.

