## Supplementary material for "Genome-wide identification of the oat *MLO* family and identification of a candidate *AsMLO* associated with powdery mildew susceptibility": Figure S1

### Slide 1
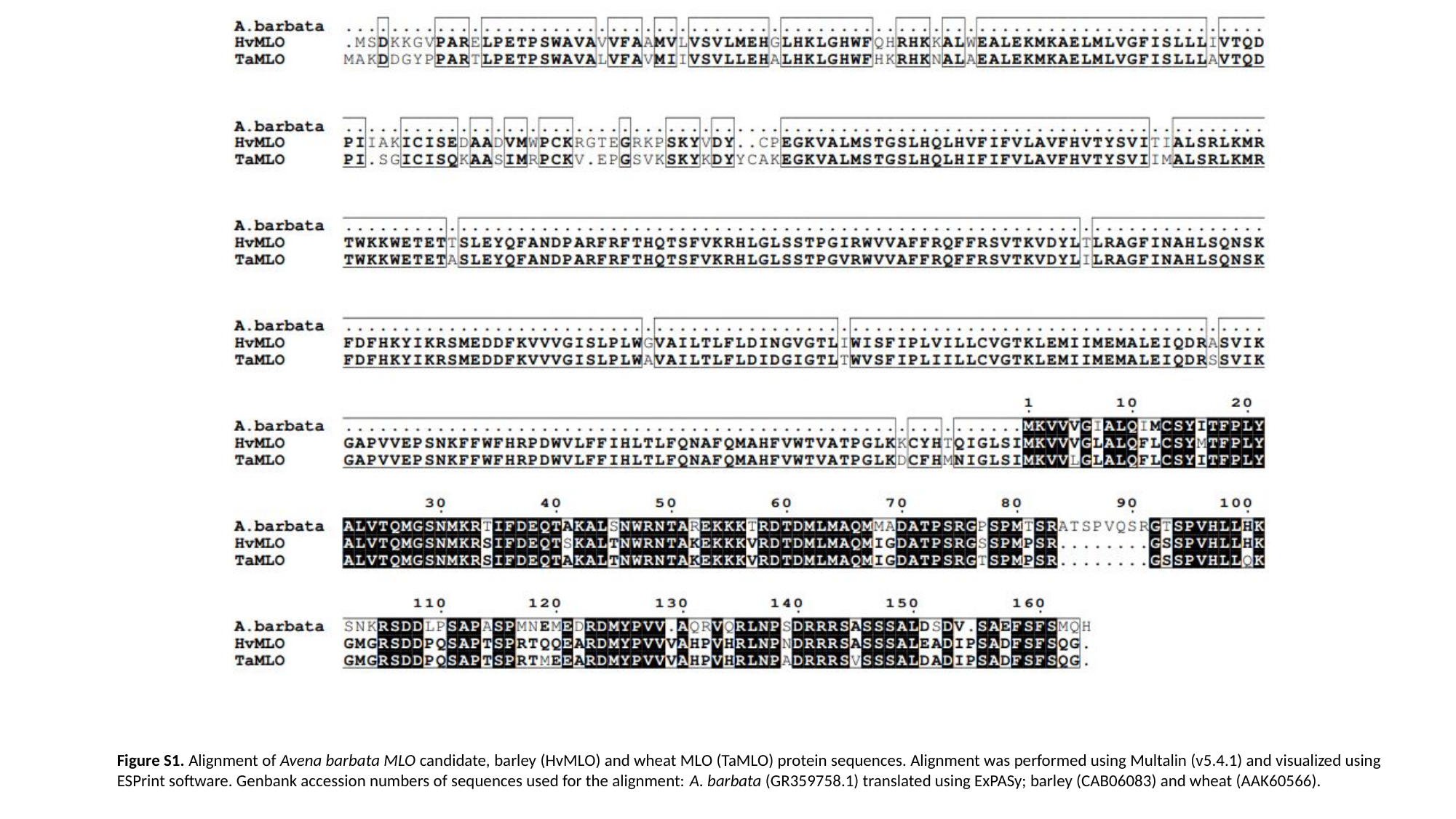

Figure S1. Alignment of Avena barbata MLO candidate, barley (HvMLO) and wheat MLO (TaMLO) protein sequences. Alignment was performed using Multalin (v5.4.1) and visualized using ESPrint software. Genbank accession numbers of sequences used for the alignment: A. barbata (GR359758.1) translated using ExPASy; barley (CAB06083) and wheat (AAK60566).
