## Supplementary material for "Genome-wide identification of the oat *MLO* family and identification of a candidate *AsMLO* associated with powdery mildew susceptibility": Figure S2

### Slide 1
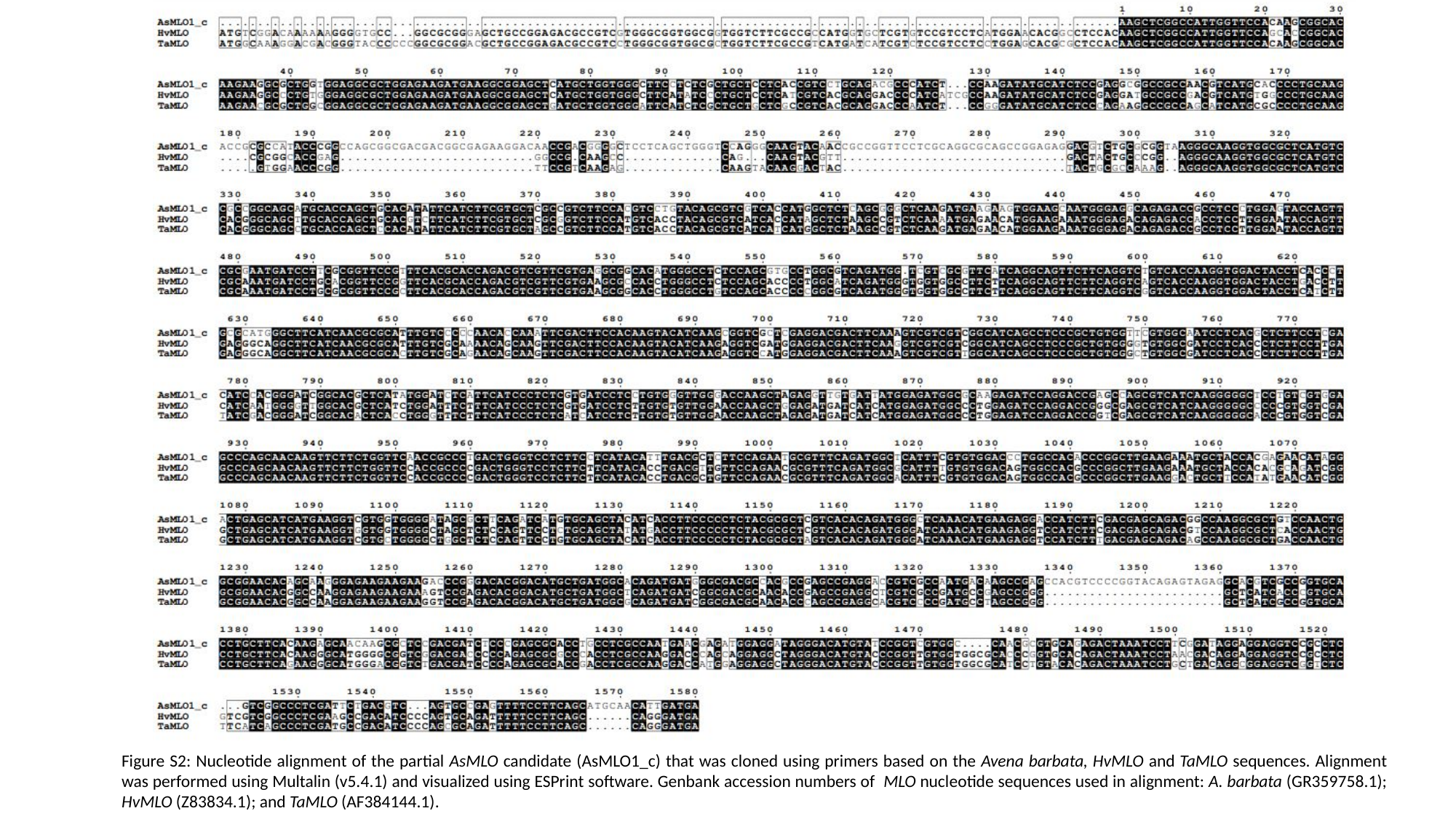

Figure S2: Nucleotide alignment of the partial AsMLO candidate (AsMLO1_c) that was cloned using primers based on the Avena barbata, HvMLO and TaMLO sequences. Alignment was performed using Multalin (v5.4.1) and visualized using ESPrint software. Genbank accession numbers of MLO nucleotide sequences used in alignment: A. barbata (GR359758.1); HvMLO (Z83834.1); and TaMLO (AF384144.1).
