## Supplementary figures and images for "Genome-wide identification of the oat *MLO* family and identification of a candidate *AsMLO* associated with powdery mildew susceptibility"

### Figure S3

#### Slide 1
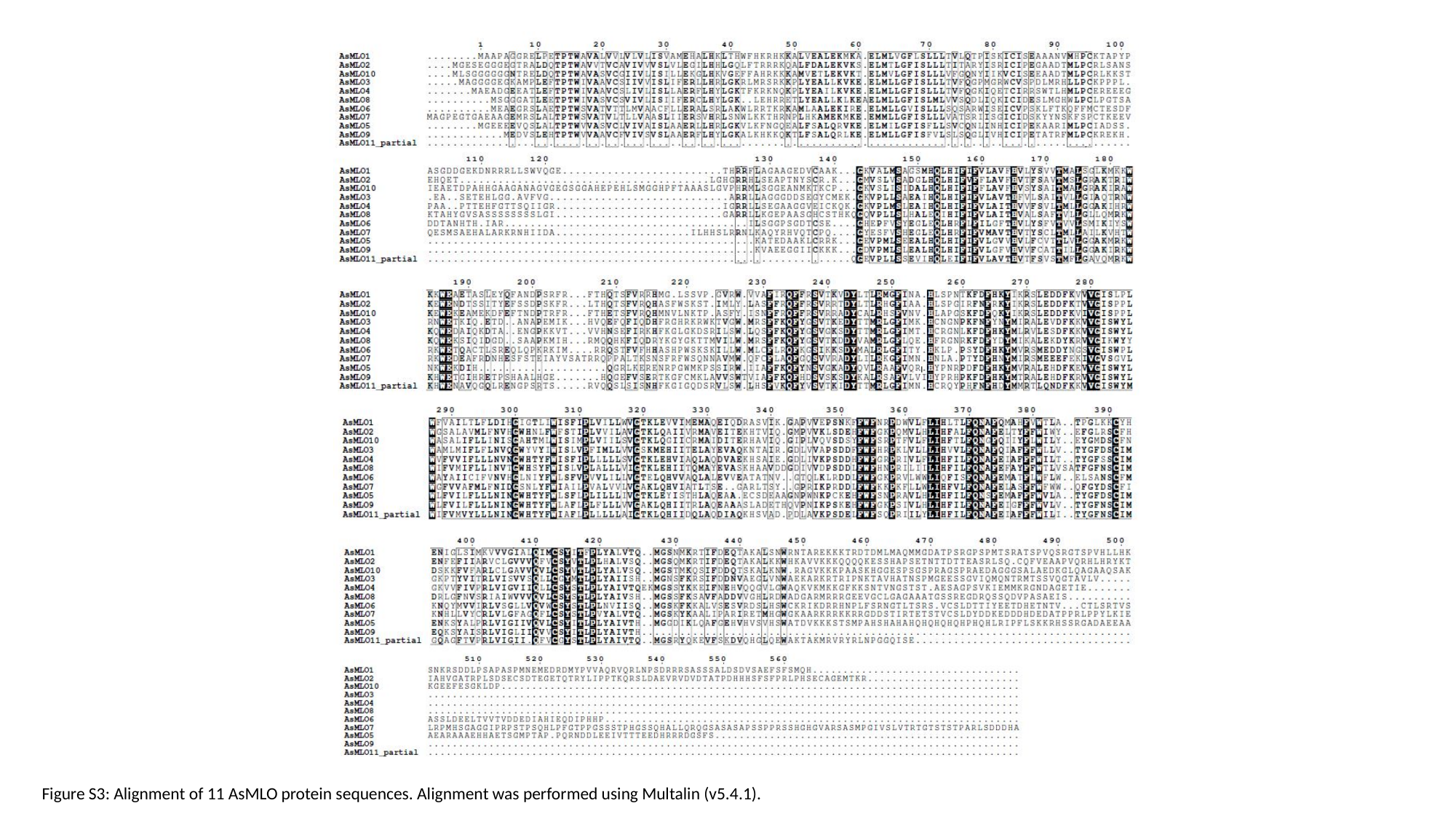

Figure S3: Alignment of 11 AsMLO protein sequences. Alignment was performed using Multalin (v5.4.1).
