## Supplementary material for "Genome-wide identification of the oat *MLO* family and identification of a candidate *AsMLO* associated with powdery mildew susceptibility": Table S1

| **Primer** | **Sequence** | **Purpose** |
| --- | --- | --- |
| As28SrRNA_F | CCTGATCTTCTGTGAAGGGTTCGA | qRT-PCR reference gene |
| As28SrRNA_R | GGTTCGATTAGTCTTTCGCCCCTA | qRT-PCR reference gene |
| AsMLO1.1_F | GTACCAGTTCGCAAATGATCC | qRT-PCR |
| AsMLO1.1_R | TCATCAGGCAGTTCTTCAGGT | qRT-PCR |
| O+B_F | GGGGACAAGTTTGTACAAAAAAGCAGGCTGCATGCTACCACGAGAACATAAGGT | *A. barbata* and barley primer with Gateway attachment |
| O+B_R | GGGGACCACTTTGTACAAGAAAGCTGGGTCTCATTGGCGACGGTCCTCGGC | *A. barbata* and barley primer with Gateway attachment |
| W+B_F | GGGGACAAGTTTGTACAAAAAAGCAGGCTGCCTGCCGGAGACGCCGTCGTGG | Wheat and barley primer with Gateway attachment |
| W+B_R | GGGGACCACTTTGTACAAGAAAGCTGGGTCCTAGCTAAAGGAAAACTCGGCACT | Wheat and barley primer with Gateway attachment |
| Uni_F | GGGGACAAGTTTGTACAAAAAAGCAGGCTGCGTCACCAAGGTGGACTACCT | Primer used for sequencing initial *AsMLO* candidate |
| Uni_R | GGGGACCACTTTGTACAAGAAAGCTGGGTCGTCTGCTCGTCGAAGATGG | Primer used for sequencing initial *AsMLO* candidate |

Table S1: List of primers used in this study
